# When Not to Kill Your Host in Competitive Bust-Boom Environments

**DOI:** 10.1101/2025.02.24.639854

**Authors:** Oskar Struer Lund, Kim Sneppen

## Abstract

Temperate phages, which can either kill their host cells or integrate into them, struggle to compete with purely virulent phages in environments with plenty of available hosts. This suggests that their survival strategy is fine-tuned for unstable conditions, where they hedge their bets between immediate replication and long-term persistence as an integrated prophage. In this study, we explored how temperate phages make these life-or-death decisions, both in isolation and when competing with other phages. We found that when temperate phages compete with each other, those with relatively stable lysogens survive better. Environments with competitive temperate phages further select for lower lysogeny frequency. Our findings suggest that dosage-dependent lysogeny choice is adapted to competing phages with overlapping immunity. In environments where phages can disperse between separate sub-systems that fluctuate independently, temperate phages struggle to survive against virulent phages.

## INTRODUCTION

Bet-hedging is a widely observed survival strategy in nature, where individuals or populations balance potential gains against the risk of loss in uncertain environments. This approach is particularly advantageous in rapidly changing conditions. When individuals choose diverse responses the population increases its chances of long-term survival. This strategic variability enables populations to persist despite environmental fluctuations. Notable examples include seed dormancy in plants to withstand unpredictable moisture levels [1], microbial stress responses that enhance survival in fluctuating conditions [2, 3], and the lysis-lysogeny decision in temperate phages, which balances immediate replication with long-term persistence [4]. However, when hedging bets, one must choose whether to prioritize long-term growth or long-term survival. Ultimately, the outcome also depends on the risk exposure for the hedged subpopulation. Here we will explore optimal bet-hedging for temperate phages with and without competing phages.

Temperate phages present one of nature’s most iconic developmental choices, the choice between the lytic amplification with the death of its host and lysogenic dormancy with coexistence. The study of this choice using phage *λ* as a model organism [5] has taught us numerous lessons about gene regulation [6, 7], epigenetics [8], developmental pathways [9–12] and regulatory landscapes [13– 15] including spontaneous escape from lysogeny with a rate *σ*∼ 10^*−*5^ per generation for the UV-inducible phage *λ* [16–18] and *σ*∼ 10^*−*4^ to 10^*−*5^ for the non-inducible phage P2 [19]. The central ability to choose between the life and death of its host implies that the phage addresses the question of an optimal lysogeny frequency *x* upon infection (varies between 0.005 and 0.5 for phage *λ* [12] dependent on number of co-infecting phages). The answer is context-dependent, and the phage can only address it using the limited information available to the phage inside its host [14, 20–22]. The lysogenic pathway comes with the price of a lower growth rate, compared to the lytic pathway [23]. It has been argued that this disadvantage is overcome by the ability of prophages to survive at very low host densities [24], while high host abundance favors virulence.

Most strikingly, temperate phages carry advantages associated with varying environments, where lytic growth is episodically inhibited by periods of low host density [4, 23]. Following Kelly’s work [25], Maslov et al. [4] explored a game-theoretical framework for growth in environments that fluctuate between “bad” and “good” periods. At each timestep, the environment is classified as bad with probability *p* or good otherwise. During bad periods the host density is low and all free phages die and only lysogens survive. Good periods support lytic growth with an amplification factor Ω relative to growth of lysogens. Assuming phages can dynamically reallocate a fraction *x* of their total population into lysogeny at each timestep, optimizing long-term growth predicts that [4]

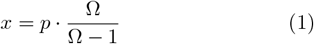

This equation optimizes the typical long-time exponential growth in an unlimited system. Realistically, the “bad events” mentioned above should represent extended “bad periods” where the prophage and its host persistently live far from other potential hosts. We will allow these bad periods to be distributed exponentially around a mean duration *τ* = 10^3^ generations reflecting a microbial world with extended periods of low-density hosts. We will see that the mean duration *τ* has marginal value influences the optimal *x* value. However, a large *τ* will predict that prophages should have a small escape probability from lysogens per generation *σ <<* 1 which is in agreement with known measurements on temperate phages [16–19].

After a bad period, no free phages are available. Hence *σ* is seeding a new free phage population at the beginning of a good period. Thus, the growth argument leading to eq. 1 should be generalized to include that only a fraction *σ* of lysogens would be available as free phages in each generation, see Fig. 1. This refinement of eq. 1 is defined in the algorithm outlined in eq. 2 in the method section and examined in Table I. Overall it predicts an optimal lysogen fraction *x* ∼ 0.5 · *p* and an optimal prophage stability per cell generation *σ* below one divided by the average length of a bad period *τ*. Thus securing long-term storage in lysogens reduces the need for bet-hedging compared to eq. 1, and reduce it more for larger values of *τ*. Importantly, these predictions do not include the influence of competing phages, i.e. that lysogeny would be disfavoured if another phage species co-infect a host, and potentially eliminate its resident prophages. Temperate phages contain defense systems against other phages [26– 30], implying that their survival depends on adaptation to competitor phages. Also, there exist phages that actively change both *x* [31] and *σ* [22] as a function of the density of nearby lysogens. These perspectives motivate us to examine the optimal lysogeny frequency *x* and stability *σ* in the context of competition with other temperate or virulent phages.

**FIG. 1.**
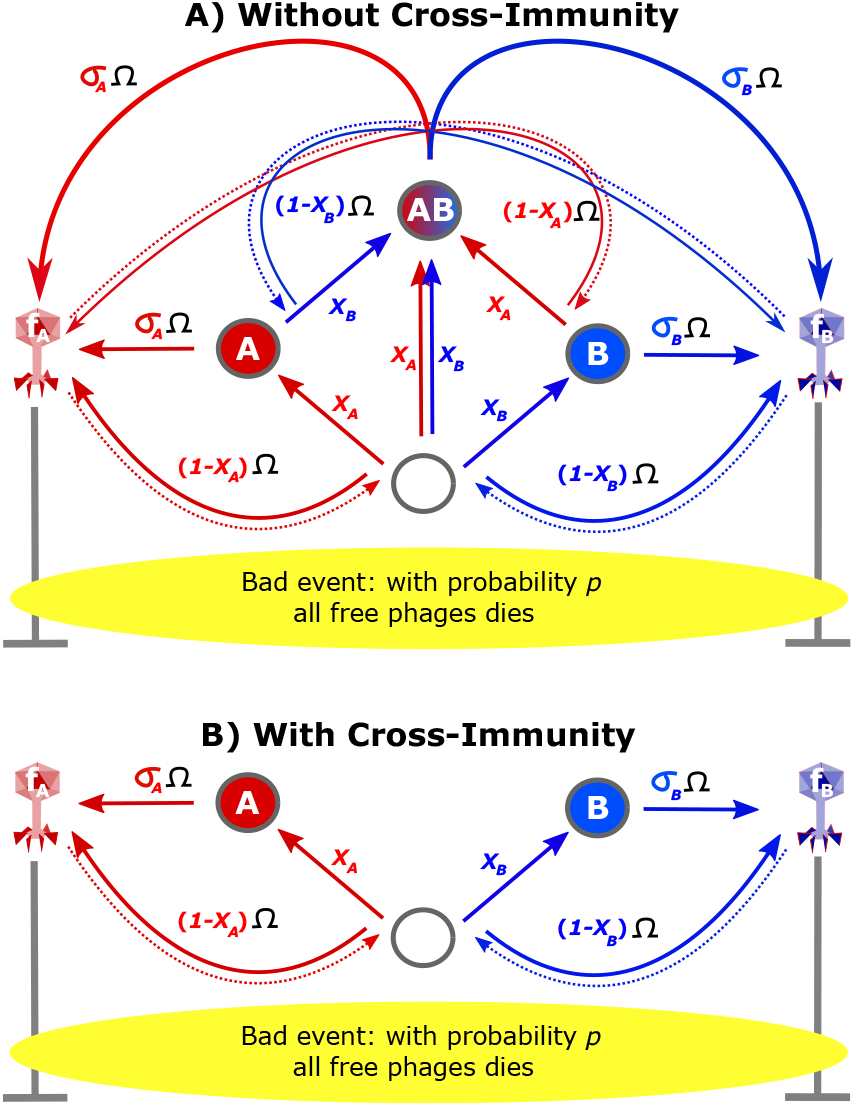
Model for competing lysogens: The figure illustrates the setup where two temperate phages with parameters (*x*_*A*_, *σ*_*A*_) and (*x*_*B*_, *σ*_*B*_), compete for a population of bacteria. The circles refer to bacteria with no prophages (empty circles with density *ρ*_0_), lysogens (A or B), or double lysogens (AB). The model is simulated in events mimicking bust-boom dynamics, with probability *p* for bad events of typical duration *τ* = 10^3^, where all free phages die (indicated by yellow). Subsequently, prophages lose populations exponentially with the duration of the bad event (see eq. 3). Otherwise, free phages may infect and potentially lyse non-immune hosts in what we denote as good events. *σ*_*A*_ and *σ*_*B*_ parameterize the stability of lysogens and Ω is the maximum reproduction number of lytic development during a good period. In our simulations this is capped by a maximum bacterial population size of 1, indicating a limited system. Panel A) shows the case where two unrelated temperate phages compete, with the possibility of forming double lysogens. This is done through eqs. 3, 4, 5 and 6. Panel B) illustrates the case where the phage has cross-immunity, simplifying the possible states. This model uses eqs: 3, 4, 6 and 7.

## RESULTS

Importantly, most of our analysis optimizes for survival quantified by mutual exclusion, and not exponential growth as was done in the Kelly bet-hedging strategy.

Fig. 1 visualizes our model for two competing phages, with arrows indicating the flow of phages through the different states. The flows between bacteria populations are associated with lysogen probability *x*_*A*_ and *x*_*B*_. Arrows from bacteria to phages mark spontaneous lysis events with frequencies *σ*_*A*_ and *σ*_*B*_. The model is updated in steps, that are “bad” with probability *p*, or otherwise good. At each step, some new uninfected bacteria are introduced to the limited environment as described in detail in the method section. Fig. 1 a) allows phages to invade each other’s lysogens, while panel b) defines competition between phages with mutual immunity.

Figure 2 a,b) illustrates the competition of two temperate phages without cross-immunity (Fig. 1 a), with different *x*_*A*_ *< x*_*B*_ = *x*(*Kelly*) = 8%, but with the same lysogen stability *σ*. Phage B in Fig. 2 a,b) have parameters that optimize long-term growth predicted by eq. 2. One observes the dominance of the phage with the smaller value of *x*, illustrating that the optimal lysogeny strategy is substantially below the Kelly optimum value of *x*(*Kelly*) ∼ 8%. Quantifying the competition, we find that the *x*_*A*_ = 2% phage excludes other phages in 99.7% of our simulations.

**FIG. 2.**
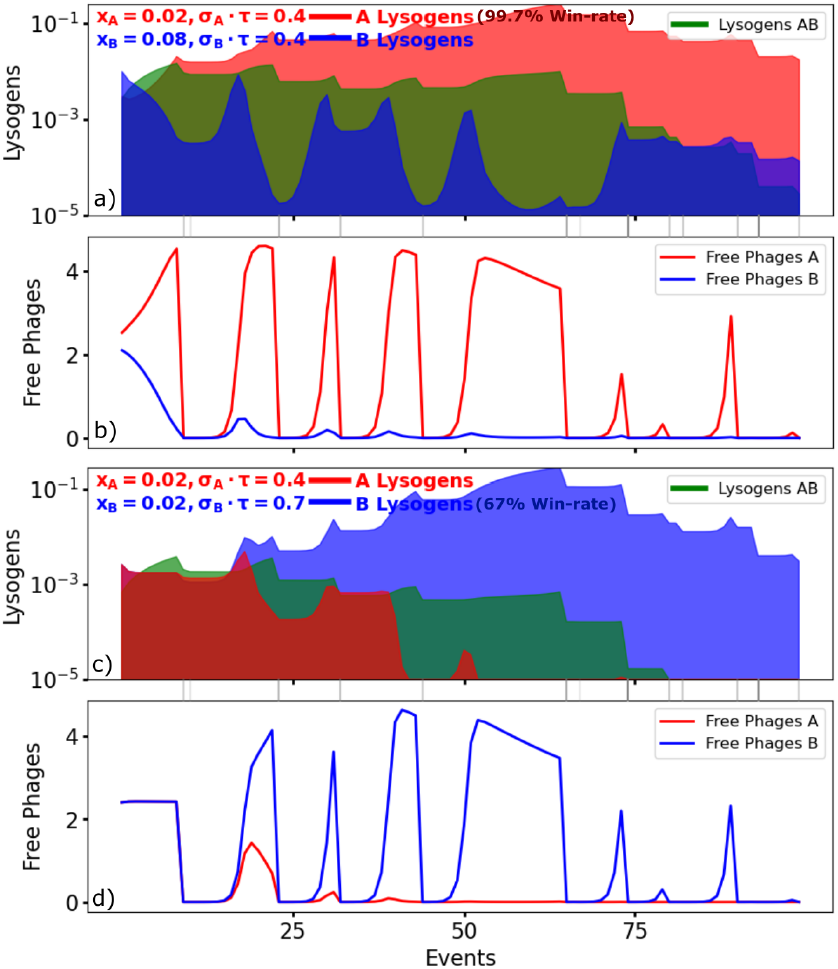
Trajectory of phage and prophage (lysogens) populations with update number, where each bad event is counted as one update. The bad events are marked with gray vertical lines between panels, with darkness their duration (*t*_*bad*_). The environmental parameters are *p* = 0.1 and the maximal lytic growth is Ω = 5. Panels a, and b) show the competition between two phages where B has *x*_*B*_ at the Kelly optimum of eq. 2. One notice that this is dominated by the more lytic A with *x*_*A*_ *< x*_*B*_, a larger chance to the exclusion of its opponent. Strategy A excludes B in 99.7% of the time series. Panels c,d) compare strategies with different exit rates from lysogeny *σ*_*B*_ *> σ*_*A*_, but with the same entry probability *x* = 0.02. Strategy B) excludes strategy A in 67% of the time series.

Fig. 2 c,d) compare phages with different *σ* values, with the same sequence of bad events as in the upper panels. We notice that a less stable lysogen outperforms the phage with a *σ* at the Kelly optimal value *σ* ∼ 0.4*/τ*. Fig. 3 systematize this competition, quantifying the success of a phage in terms of the chance to exclude a competitor. All panels have common parameters *p* = 0.1, Ω = 5. Fig. 3 a) explores the optimal *x, σ* systematically in competitions where the phages have different immunity (Fig. 1a), by plotting the probability of excluding the optimal strategy (*x, σ*) = (2, 0%, 0.8*/τ*) as a function of phage parameters. The figure highlights that the two-phage competition favors lower lysogeny frequency but similar stability than predicted from the non-competitive Kelly-like situation 2. The low optimal *x* value reflects that lysogens are exposed to predation by the other phage.

**FIG. 3.**
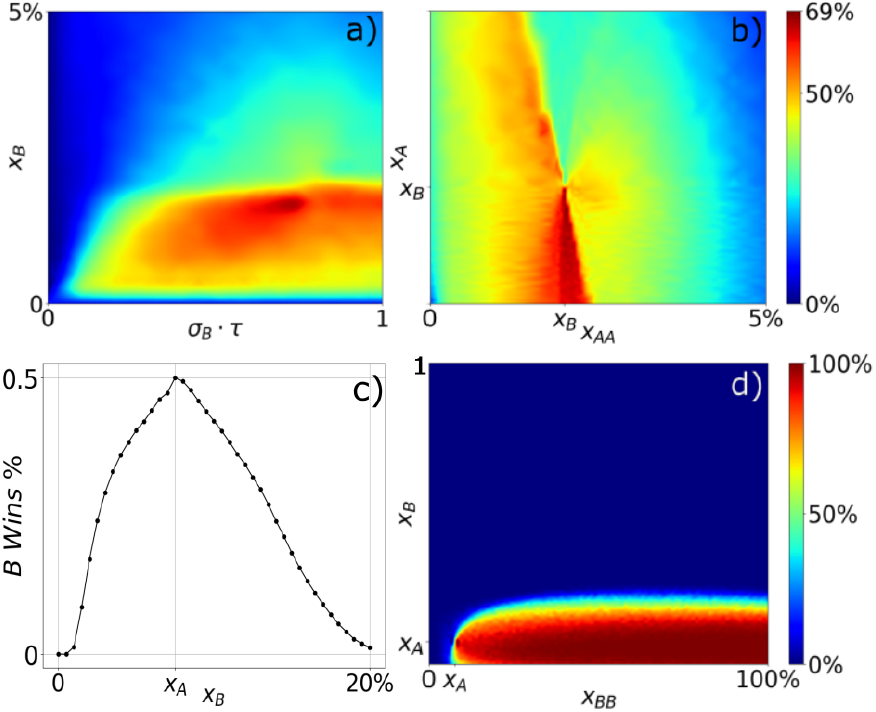
All four plots have common *p* = 0.1, Ω = 5. Panels and b) assumes no cross-immunity. a) shows optimal *x, σ*choices, indicating that optimal lysogeny frequency is *x* = 2%, paired with a lysogeny stability of *σ · τ* = 0.8. Color marks the probability that the optimum excludes competitors with properties given by coordinates. The optimal strategy found in a) is then tested in panel b) against a MOI-sensitive phage. Here we find that multiple infections *x*_*AA*_ *∼* 2% is optimal with single infection optimal lysogeny frequencies, *x*_*A*_, ranging from 0 to 1.9%. Plot c,d) simulate cross-immunity see Fig. 1b). Plot c) optimizes lysogeny frequency with fixed *σ* from growth optimization, see eq. 2 and table Ia, obtaining a maximum at *x*_*A*_ = 7.5%. d) compare MOI phages B against the optimum from c). The plot shows that when *x*_*B*_ *< x*_*A*_ and *x*_*BB*_ *> x*_*A*_, the *MOI*-phage, *B*, is superior to *A*, with lysogeny frequency *x*_*A*_.

Fig. 3 b) extends the lysogeny choices, by allowing the infecting phage to sense whether it is alone in its infected host, or has co-infected with another phage of the same type. In the *λ* phage this counting is facilitated by the regulatory protein CII [9, 14, 32, 33]. Phages with this dosage dependence are denoted MOI-sensitive. Panel b) considers the competition between this *λ* like phage and another dose-independent (P2-like) phage assigned the optimal value of *x, σ* from Fig. 3.

Fig. 3c) optimizes for optimal lysogeny frequency with mutual immunity (using the model from Fig 1b), with fixed *σ* · *τ* = 0.4, mirroring *Kelly optimal σ*. Simulation shows that *x* = 7.5% is optimal, marked with “*x*_*A*_”. Fig. 3d) takes the optimal strategy from panel and competes it with a phage that can differentiate *x* between single and multiple infections (MOI sensitive phage). Here, we find that any *x*_*B*_ *<* 7.5% combined with any *x*_*BB*_ *>* 7.5% beats the phage without the ability to sense double infections in the vast majority of simulations.

Table I.B) summarize the optimal *x, σ* in different environments, varying *p* and Ω. At low Ω the optimal *x* is substantially below the optimal values from growth optimization (eq. 2). Especially at *p* = 0.1 the optimal *x* ∼ 2% is a factor 4 below the optimal of 8% from I.A). At higher Ω the situation is more unclear, with an optimum lysogen frequency exhibiting a cyclical context dependence on who out-competes who. At high Ω, the death of one, but not the other phage, is associated with particular sequences of events. This is explored in Fig. S1, where the bottom panel quantifies the last 40 timesteps in sequences where one phage survives the other: The cycles reflect that a low *x* ∼ 1% wins over a very high *x*, while it can be replaced in sequentially increasing *x* competitions because of their dominance in time sequences with unusually many bad events. However, when considering a phage with high *x* it becomes fragile to a near lytic strategy (*x* small).

Table IC) examines optimal strategies when exposed to phages with shared immunity. This corresponds to situations that would be common during evolution, as mutations may change *x* or *σ* without changing the properties of the repressor for immunity maintenance. The model with the shared immunity is simulated as in Fig. 1B and in eq. 7. Notably, by eliminating the possibility to take over the lysogen of other phages the requirement for longtime survival favors arbitrarily small *σ* (set by threshold *ϵ*). Here we freeze the *σ* to the values from table IA) corresponding to maximal solitary growth.

Remarkably, the optimal *x* values in the cross-immunity competition in Table IC) show that they are quite close to the values found in the modified Kelly optimization (Table IA)). Table IC columns 4,5 compares the optimal single *x*_*A*_ phage with a MOI-sensitive phage.

The table shows that the single *x*_*A*_ phage is beaten by any phage where *x*_*B*_ *< x*_*A*_ and *x*_*BB*_ *> x*_*A*_, i.e. any phage with dose-sensitive lysogen choice as illustrated by the red area in Fig 3 c). This result is markedly different from MOI-sensitive phages competing against phages without cross-immunity (see Table IB). Without cross-immunity, it may even be preferable with *x*_*AA*_ *< x*_*A*_ when Ω is large.

The model allows for an extension to include that a real environment exchanges microbes with other environments. This provides the virulent phage an opportunity to re-establish itself after a bad period, thereby reducing the advantage of temperate phages. Eq. 8 and the supplement describe a formalism where each environment is assumed to weakly exchange microbes with many other similar but independently fluctuating systems. Because the duration of bad periods in each environment is much longer than good periods (i.e. that *τ >>* 1*/p*), most exchanged phages will be lost.

The mutual exclusion between a temperate phage and a virulent one is explored in Fig. 4. We see that temperate phages are only competitive when the flux between the environments is smaller than a threshold set by *σ* and *p*. Panel b) explores the MOI sensitive *x*_*A*_, *x*_*AA*_ with fixed *x*_*A*_ = 0, and *x* = *x*_*AA*_ corresponding to shown color. The corresponding plot with fixed *x*_*A*_ = 10^*−*3^ is in supplement Fig. S3. This suggests that MOI sensitivity does not help against lytic phages. Panel c) shows that a higher probability of bad environments greatly benefits temperate phages as also predicted in [4]. In all panels a-c) one observes that high effective burst size favors an increasingly lytic behavior.

**FIG. 4.**
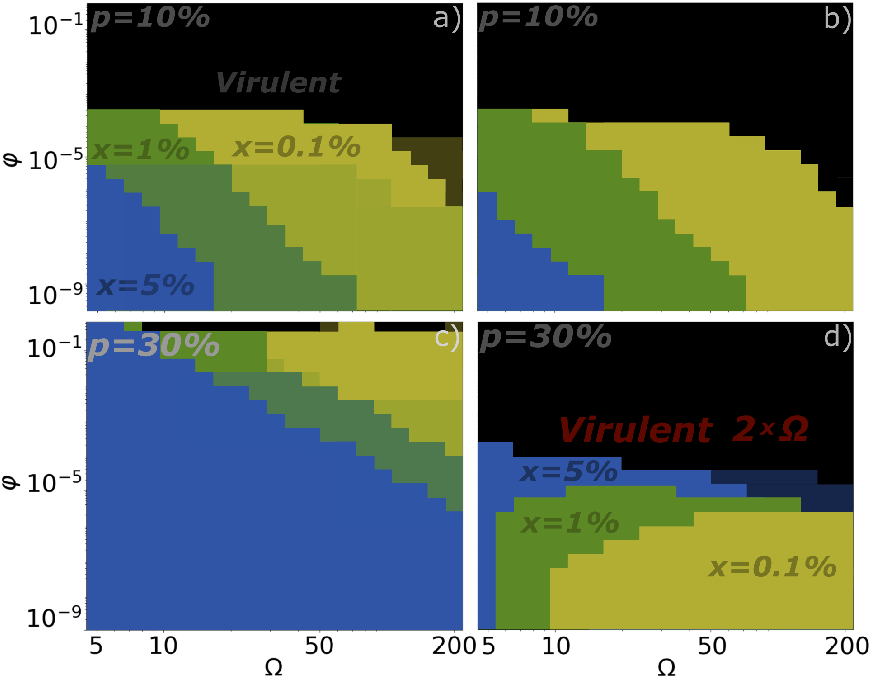
Competition between a temperate phage and a virulent phage for coupled environments with different burstsizes, Ω, and fluxes, *φ*, between the environments. The model outlined in eq. 8. **a)** Environments with the frequency of bad events *p* = 10%. The plot is divided into four regions, each marked with the winning phage type. The black region represents conditions where virulent phages are superior to any temperate strategy. The yellow, green, and blue indicate parameters when a temperate strategy with *x* = 0.1% (yellow),*x* = 1% (green), respectively *x* = 5% (blue) wins over the virulent phage. Shaded regions mean neither the virulent nor the temperate phage dies within 10^10^ events. **b)** Dosesensitive (MOI) phages tested against a virulent phage. The MOI phage is lytic if single infectious, but has a lysogeny frequency for multiple infections. The colors refer to lysogeny frequencies for multiple infections. There are no shaded areas, indicating that a phage is always excluded within 10^10^ events. **c)** As in a) but with *p* = 30%. **d)** as in c) but with the virulent phage having twice the burst size of its competing temperate phage. Here larger lysogeny frequencies become more dominant for the competitive fluxes.

Panels a-c) use the same Ω for lytic and temperate phages, while real temperate phages on average have about 75% longer latency than virulent phages (averages calculated from ref. [34]). The ability to increase replication naturally favors the purely virulent phages more, as seen by comparing panels d) and c). Interestingly, when competing with a more productive lytic phage, the temperate phage does best when it deviates strongly from virulence, i.e. has a high *x*.

## DISCUSSION

Bacteria capable of harboring prophages [35] often exhibit faster growth in nutrient-rich environments, suggesting that phage-bacterial dynamics are closely linked to ecological conditions. While all bacteria face periods of nutrient scarcity, those that thrive in fluctuating environments tend to carry phages employing bet-hedging strategies to endure harsh conditions. In contrast, bacteria that experience relatively stable growth across different conditions are more likely to coexist with predominantly virulent phages. For example, oceanic cyanobacteria, whose growth is more steady from generation to generation, are almost exclusively infected by virulent phages [36].

The coexistence of virulent and temperate phages has been explored under constant [37] and fluctuating [4, 23] environmental conditions. Theoretical work by [37] suggests that temperate phages can invade a lytic population in periodically diluted environments, where lysogens outcompete free phages in the long run. This aligns with scenarios where the free phage decay rate significantly exceeds the infection rate. Additionally, [38] showed that in constant environments, optimal lysogen stability *σ* should be minimized, constrained only by the growth disadvantage of lysogens. Our paper, in contrast, explores temperate phages in the context of variable environments. In fluctuating environments, we predict that a finite *σ* is favored because it is needed to seed new free phages in the beginning of good periods.

Our findings extend the Kelly bet-hedging framework [4, 25] to account for phage competition in boom-bust conditions with short periods of huge returns and very long periods where survival is challenging. These extreme conditions can be deduced from the many temperate phages that typically have a large probability to “bet” on the lytic development (1− *x*∼ 1) upon infection, complemented with a low rate *σ* ∼ 10^*−*4^ of leaving the more secure existence as a prophage. This leaves temperate phages as a compromise of growth with survival during long bad periods. However, their existence as an integrated prophage leaves them vulnerable to other phages that may kill or steal their relatively safe “investment” in the otherwise durable lysogen.

The subsequent model in our paper teaches us that competition from widely different temperate phages can make the optimal lysogeny fraction *x* substantially below the value predicted from optimizing long-term growth without competition. The second lesson is that the possible elimination of a lysogen by competitors only has a moderate effect on the optimal stability of lysogeny, *σ*. This reinforces the prediction that an experimentally reported value of *σ* ∼ 10^*−*4^ to 10^*−*5^ [17–19] reflect that individual bacterial strains often persist in low abundance during long periods in the wild.

When competing temperate phages that are mutually immune (Table 1C), the optimal values of *x* remain close to that from Table 1A maximizing long-term growth. The higher *x* reflects that in the immune case, the lysogens are safe from infection by competing phages. Thus mutually immune phages do not drive each other to increased virulence either in the form of smaller *x* or larger *σ*. The role of the multiplicity of infection (MOI) is particularly significant in competition between closely related phages. Studies have shown that phage *λ* increases its lysogeny frequency by nearly 100-fold under double infection [9, 12]. Our mutual immunity model confirms that phages with MOI-sensitive lysogeny strategies consistently outcompete those with fixed lysogeny probabilities, suggesting an evolutionary advantage for dosage-dependent decision-making. In this context, interestingly, *λ* phages exhibit MOI-sensitive lysogeny, while P2like phages are dosage-independent, reflecting divergent evolutionary pressures.

**TABLE 1.**
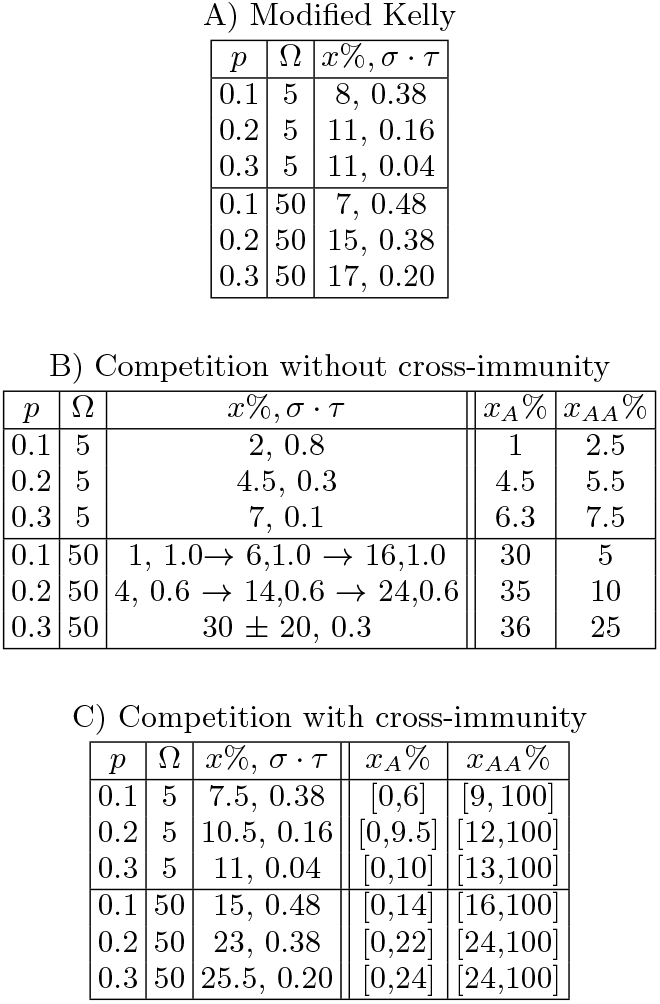
Optimal lysogenization frequencies with environmental parameter *τ* = 1000. Panel A) shows the “welltemperate-phage” from optimizing eq. 2 numerically. For *τ* = 10 and Ω = 50 the optima are (9%,0.6), (17%,0.5) and (26%,0.4) for *p* = 0.1,0.2, and 0.3 respectively. B) Optimal competitor against phages without cross-immunity (Fig. 1a). Low *p* and high Ω are associated with context-dependent winners in a “rock-paper-scissor” like competition, see Fig S1. For *p* = 0.1 and *p* = 0.2, the arrows point to a winning strategy given competition against the *x, σ* at the arrow’s base. The *x*_*A*_ and *x*_*AA*_ columns list optimal MOI-sensitive phage competing with the optimum *x* in column 4, keeping *σ* fixed. C) Optimal competitors with cross-immunity (Fig. 1b), maintaining *σ* from table A. Phages with one *x* value will be out-competed by MOI-sensitive phages, provided that *x*_*A*_ is anywhere below *x* and *x*_*AA*_ anywhere above *x*.

| $p$ | $\Omega$ | $x\%, \sigma \cdot \tau$ |
| --- | --- | --- |
| 0.1 | 5 | 8, 0.38 |
| 0.2 | 5 | 11, 0.16 |
| 0.3 | 5 | 11, 0.04 |
| 0.1 | 50 | 7, 0.48 |
| 0.2 | 50 | 15, 0.38 |
| 0.3 | 50 | 17, 0.20 |

B) Competition without cross-immunity
| $p$ | $\Omega$ | $x\%, \sigma \cdot \tau$ | $x_A\%$ | $x_{AA}\%$ |
| --- | --- | --- | --- | --- |
| 0.1 | 5 | 2, 0.8 | 1 | 2.5 |
| 0.2 | 5 | 4.5, 0.3 | 4.5 | 5.5 |
| 0.3 | 5 | 7, 0.1 | 6.3 | 7.5 |
| 0.1 | 50 | 1, 1.0 → 6, 1.0 → 16, 1.0 | 30 | 5 |
| 0.2 | 50 | 4, 0.6 → 14, 0.6 → 24, 0.6 | 35 | 10 |
| 0.3 | 50 | 30 ± 20, 0.3 | 36 | 25 |

TABLE I. Optimal lysogenization frequencies with environmental parameter $\tau = 1000$ .
| $p$ | $\Omega$ | $x\%, \sigma \cdot \tau$ | $x_A\%$ | $x_{AA}\%$ |
| --- | --- | --- | --- | --- |
| 0.1 | 5 | 7.5, 0.38 | [0,6] | [9,100] |
| 0.2 | 5 | 10.5, 0.16 | [0,9.5] | [12,100] |
| 0.3 | 5 | 11, 0.04 | [0,10] | [13,100] |
| 0.1 | 50 | 15, 0.48 | [0,14] | [16,100] |
| 0.2 | 50 | 23, 0.38 | [0,22] | [24,100] |
| 0.3 | 50 | 25.5, 0.20 | [0,24] | [24,100] |

Finally, our model allows us to address the influence of coupled independent environments. The overall prediction is that a high enough transfer (*φ* large) serving the virulent phage with the bet-hedging of multiple environments, can make them exclude temperate phages. This aligns with empirical observations: temperate phages dominate in the gut microbiome, where environmental fluctuations are relatively contained [39], whereas oceanic cyanobacteria, subject to high mixing, carry few prophages [36].

## MODELS

The main analysis of the paper is to find optimal *x, σ* for different values of the phage and environmental parameters Ω, *p*, with results shown in table 1.

For comparison, we first extend the “Well-temperate phage” scenario [4] to lysogens as long-term storage, that only are partially available for eventual lysis at each timestep. In this simple extension, we first consider one type of phage, subdivided into a free phage population *f*, and prophage population *ρ* bound in lysogens. The transition between these states is set by parameters *x* defining lysogenization probability for the free phage meeting a host, and *σ* determining the spontaneous lysis rate of lysogens.

The model is executed in discrete time steps, which are assigned to be either bad or good. This mimics a burst-and-bust view of bacterial existence, where relatively short periods with abundant resources and potentially plenty of bacterial hosts are interrupted by bad periods where bacterial density is so low that the phage cannot find them. At every timestep, the probability of entering a bad period has probability *p*. The length of the bad period is expected to be longer than the good periods, parameterized by a characteristic length *τ* timestep. When a population of phages and bacteria enters a bad period the free phages are lost, while the lysogens lyse with some phage-dependent rate *σ*. If the bacteria does not enter the bad period it is in a good (burst) period, where free phages proliferate or enter lysogens as described by a phage-dependent lysogenization rate *x*. If phages have effective bust size Ω one update step is defined by:

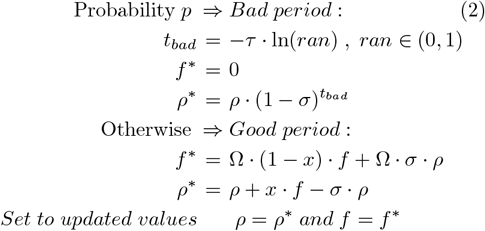

In this Kelly-like scenario, we optimize the long-term exponential growth of the total phage genomes, *f* + *ρ*.

The main model of this paper describes a biological system with two temperate phages that compete with each other as outlined in Fig. 1A). The basic description of the boom-bust dynamics is as in the above model, but with a limited host population. This prevents us from estimating long-term exponential growth but allows us to include competition between the two phages with basic parameters (*x*_*A*_, *σ*_*A*_) and (*x*_*B*_, *σ*_*B*_). In our extension of eq. 2 we assume that all the available phages of a given type *f*_*x*_ should be distributed randomly between all available host *ρ* = *ρ*_0_ + *ρ*_*A*_ + *ρ*_*B*_ + *ρ*_*AB*_. Counting these densities in discrete numbers, the probability that a given bacteria is not infected by one phage is 1 −1*/ρ*. The probability that a given bacteria is infected by one of *f*_*x*_ phages is 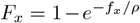. This equation only deals with ratios of numbers, that can be rescaled to densities. One update step is now defined by:

**Bad times chosen with probability** *p*:

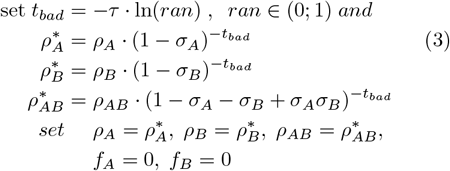

**Good times chosen if bad not selected**

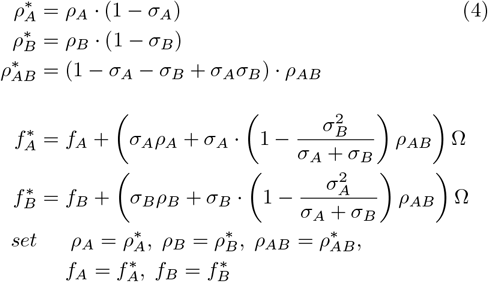

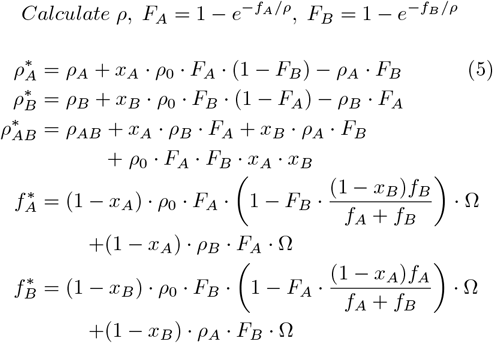

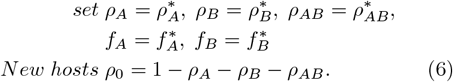

Here, the double infections of an uninfected bacteria *ρ*_0_ only leads to lysogens if both phages choose this option, while lytic descendant is set by the first infecting phage (estimated by ratios of free phages). We will optimize parameters *x*_*A*_ and *σ*_*A*_ for the lowest relative extinction risk given environmental conditions; *p, t*_*bad*_, Ω and in principle, see Table S1. When comparing two phages with different parameters, we simulate many sequences, each starting with *ρ*_0_ = 1 and *f*_*A*_ = *f*_*B*_ = 1.

For each case, we simulate until one of the phages, say phage *A* has total genome count *f*_*A*_ + *ρ*_*A*_ + *ρ*_*AB*_ below the threshold *ϵ* = 10^*−*9^. The winning phage is then the one with the largest number of survivals. Finding the surviving phage is typically done in ∼ 10^3^ steps for Ω = 5 and ∼ 10^4^ for Ω = 50. We tested that all reported results remain unchanged with different cut-off thresholds of *ϵ* = 10^*−*6^ and *ϵ* = 10^*−*12^. We never obtained co-existence.

In contrast to the maximize growth condition for the “Kelly-like” game in eq. 2, the finite environments above optimize for robustness against competitors.

Ending each update with the addition of new hosts given by eq. 6 means that phages are selected for their ability to infect new hosts. An influx of new susceptible hosts considered in the constant environment models [22, 24] where it was suggested to favor increased virulence. Influx could be implemented in a bio-reactor by replacing a fraction of lysogens with the inflow of new vacant hosts, adjusted dynamically to keep the total concentration of all bacteria constant. Bad periods might be implemented by adding appropriate salts to kill free phages. In the real world, the supply of new hosts emulates an ongoing selection pressure on phage infection properties.

Eqs. 5 assume that all phages infect bacteria at each update step. A more detailed model include a phage death term *δ* that is defined relative to the carrying capacity set to 1. This death term is implemented by changing 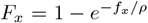 to 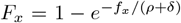 in eq. 5 for both *x* = *A* and *x* = *B*. For dense laboratory-like environments *δ <<* 1 it simplifies to the standard model. In contrast, *δ >>* 1 corresponds to so low bacteria density that a phage rarely finds a host before it dies. One could understand the bad times as situations where *δ*∼ ∞. In general, a larger *δ* would give a smaller effective burst size Ω*/ρ*→ Ω*/*(*ρ* + *δ*). The supplement table shows that simulations with *δ* = 10 give similar results a 10fold reduction in Ω in our standard model. The above model can be modified to compare phages that provide immunity against each other (Fig. 1B). That is relevant when competing phages share part of their immunity region, specifically if they have similar repressors for lysogen maintenance [40]. Mutual immunity is obtained by fixing setting *ρ*_*AB*_ = 0 and replacing updates in eq. 5 with

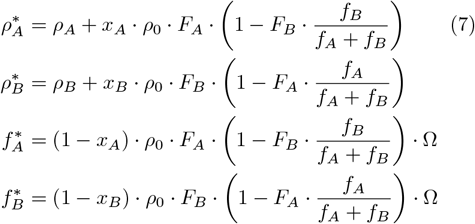

This replacement reflects that the first phage to infect a host immunizes against second infections irrespective of whether the first phage chooses lysis or lysogeny.

The above models do not consider that *x* differs between single and double infections. This can be explored by separating the probability of exactly one phage infection *P* (1) = (1 −*e*^*−f/ρ*^) · *e*^*−f/ρ*^ from the likelihood of two or more phages infecting a given bacterium *P* (≥ 2) = 1 − 2*e*^*−f/ρ*^ + *e*^*−*2*f/ρ*^. This equations are more complicated and showed in the supplement.

The above model considers only one coherent isolated environment and does not allow this to receive or transmit phages to other independent environments. We also consider the case where the single environment is coupled to an infinity of independent environments. Loss of phages from the given environment is included at the end of each time-step step by subtracting *φ*· *f*_*x*_ from phage populations. The input of phages in the given environment happens at the beginning of a time-step by adding *φ* ⟨ *f*_*x*_⟩ being the average of transmissions phage *x*. We here use the approximation that the averaged phage over many environments can be obtained from the average over the previous history:

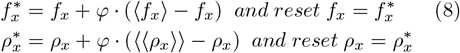

The supplement describes an approximate method for obtaining ⟨*f*⟩ *f* ⟨ (*good times*) ⟩ */pτ* and ⟨⟨*ρ*_*x*_⟩⟩ using the history of past events (See supplement “Additional algorithms”, subsection B). The supplement also explores the relative importance of the fluxes of free phages and lysogens (See Fig. S2), and contains detailed derivations of how to estimate averages of bacterial densities across many good and bad periods ⟨⟨*ρ*_*x*_⟩⟩, *x* = *A, B*.

## Supporting information

Deatils of equations, suplementary figures

## Acknowledgment

We sincerely thank Namiko Mitarai for insightful discussions and valuable feedback on the manuscript. This research was funded by the Danish National Research Foundation (grant no. DNRF170).

## References

[1] D. Cohen, Optimizing reproduction in a randomly varying environment, Journal of theoretical biology 12, 119 (1966).

[2] J.-W. Veening, W. K. Smits, and O. P. Kuipers, Bistability, epigenetics, and bet-hedging in bacteria, Annu. Rev. Microbiol. 62, 193 (2008).

[3] S. F. Levy, N. Ziv, and M. L. Siegal, Bet hedging in yeast by heterogeneous, age-correlated expression of a stress protectant, PLoS biology 10, e1001325 (2012).

[4] S. Maslov and K. Sneppen, Well-temperate phage: optimal bet-hedging against local environmental collapses, Scientific reports 5, 10523 (2015).

[5] E. M. Lederberg and J. Lederberg, Genetic studies of lysogenicity in escherichia coli, Genetics 38, 51 (1953).

[6] G. K. Ackers, A. D. Johnson, and M. A. Shea, Quantitative model for gene regulation by lambda phage repressor., Proceedings of the national academy of sciences 79, 1129 (1982).

[7] S. Roy, S. Garges, and S. Adhya, Activation and repression of transcription by differential contact: two sides of a coin, Journal of Biological Chemistry 273, 14059 (1998).

[8] M. Ptashne, A. Jeffrey, A. D. Johnson, R. Maurer, B. J. Meyer, C. O. Pabo, T. M. Roberts, and R. T. Sauer, How the lambda repressor and cro work, Cell 19, 1 (1980).

[9] P. Kourilsky and A. Knapp, Lysogenization by bacteriophage lambda: Iii.-multiplicity dependent phenomena occuring upon infection by lambda, Biochimie 56, 1517 (1975).

[10] L. Zeng, S. O. Skinner, C. Zong, J. Sippy, M. Feiss, and I. Golding, Decision making at a subcellular level determines the outcome of bacteriophage infection, Cell 141, 682 (2010).

[11] I. Golding, S. Coleman, T. V. Nguyen, and T. Yao, Decision making by temperate phages, Encyclopedia of Virology 1, 5 (2019).

[12] Y. Geng, T. V. P. Nguyen, E. Homaee, and I. Golding, Using bacterial population dynamics to count phages and their lysogens, Nature Communications 15, 7814 (2024).

[13] H. Eisen, P. Brachet, L. P. d. Silva, and F. Jacob, Regulation of repressor expression in λ, Proceedings of the National Academy of Sciences 66, 855 (1970).

[14] A. B. Oppenheim, O. Kobiler, J. Stavans, D. L. Court, and S. Adhya, Switches in bacteriophage lambda development, Annu. Rev. Genet. 39, 409 (2005).

[15] S. L. Svenningsen and S. Semsey, Commitment to lysogeny is preceded by a prolonged period of sensitivity to the late lytic regulator q in bacteriophage λ, Journal of bacteriology 196, 3582 (2014).

[16] E. Aurell, S. Brown, J. Johanson, and K. Sneppen, Stability puzzles in phage λ, Physical Review E 65, 051914 (2002).

[17] K. Baek, S. Svenningsen, H. Eisen, K. Sneppen, and S. Brown, Single-cell analysis of λ immunity regulation, Journal of molecular biology 334, 363 (2003).

[18] J. W. Little and C. B. Michalowski, Stability and instability in the lysogenic state of phage lambda, Journal of bacteriology 192, 6064 (2010).

[19] E. Six, The rate of spontaneous lysis of lysogenic bacteria, Virology 7, 328 (1959).

[20] A. Trusina, K. Sneppen, I. B. Dodd, K. E. Shearwin, and J. B. Egan, Functional alignment of regulatory networks: a study of temperate phages, PLOS Computational Biology 1, e74 (2005).

[21] H. M. Doekes, G. A. Mulder, and R. Hermsen, Repeated outbreaks drive the evolution of bacteriophage communication, Elife 10, e58410 (2021).

[22] J. B. Bruce, S. Lion, A. Buckling, E. R. Westra, and S. Gandon, Regulation of prophage induction and lysogenization by phage communication systems, Current Biology 31, 5046 (2021).

[23] F. M. Stewart and B. R. Levin, The population biology of bacterial viruses: why be temperate, Theoretical population biology 26, 93 (1984).

[24] L. M. Wahl, M. I. Betti, D. W. Dick, T. Pattenden, and A. J. Puccini, Evolutionary stability of the lysis-lysogeny decision: Why be virulent?, Evolution 73, 92 (2019).

[25] J. L. Kelly, A new interpretation of information rate, the bell system technical journal 35, 917 (1956).

[26] J. J. Weigle and M. Delbrück, Mutual exclusion between an infecting phage and a carried phage, Journal of bacteriology 62, 301 (1951).

[27] B. D. Howard, Phage lambda mutants deficient in r ii exclusion, Science 158, 1588 (1967).

[28] J. Heinrich, M. Velleman, and H. Schuster, The tripartite immunity system of phages p1 and p7, FEMS microbiology reviews 17, 121 (1995).

[29] J. Nesper, J. Blaß, M. Fountoulakis, and J. Reidl, Characterization of the major control region of vibrio cholerae bacteriophage k139: immunity, exclusion, and integration, Journal of bacteriology 181, 2902 (1999).

[30] D. Refardt, Within-host competition determines reproductive success of temperate bacteriophages, The ISME journal 5, 1451 (2011).

[31] Z. Erez, I. Steinberger-Levy, M. Shamir, S. Doron, A. Stokar-Avihail, Y. Peleg, S. Melamed, A. Leavitt, A. Savidor, S. Albeck, et al., Communication between viruses guides lysis–lysogeny decisions, Nature 541, 488 (2017).

[32] A. C. Palmer, A. Ahlgren-Berg, J. B. Egan, I. B. Dodd, and K. E. Shearwin, Potent transcriptional interference by pausing of rna polymerases over a downstream promoter, Molecular cell 34, 545 (2009).

[33] M. Avlund, S. Krishna, S. Semsey, I. B. Dodd, and K. Sneppen, Minimal gene regulatory circuits for a lysis-lysogeny choice in the presence of noise, PloS one 5, e15037 (2010).

[34] M. De Paepe and F. Taddei, Viruses’ life history: towards a mechanistic basis of a trade-off between survival and reproduction among phages, PLoS biology 4, e193 (2006).

[35] M. Touchon, A. Bernheim, and E. P. Rocha, Genetic and life-history traits associated with the distribution of prophages in bacteria, The ISME journal 10, 2744 (2016).

[36] M. F. Marston and J. B. Martiny, Genomic diversification of marine cyanophages into stable ecotypes, Environmental Microbiology 18, 4240 (2016).

[37] G. Li, M. H. Cortez, J. Dushoff, and J. S. Weitz, When to be temperate: on the fitness benefits of lysis vs. lysogeny, Virus Evolution 6, veaa042 (2020).

[38] M. G. Cortes, J. Krog, and G. Balázsi, Optimality of the spontaneous prophage induction rate, Journal of theoretical biology 483, 110005 (2019).

[39] L. Avellaneda-Franco, S. Dahlman, and J. J. Barr, The gut virome and the relevance of temperate phages in human health, Frontiers in Cellular and Infection Microbiology 13, 1241058 (2023).

[40] N. G. Carlson and J. W. Little, Highly cooperative dna binding by the coliphage hk022 repressor, Journal of molecular biology 230, 1108 (1993).

