## Supplementary material for "When Not to Kill Your Host in Competitive Bust-Boom Environments": Deatils of equations, suplementary figures

### Supplemental Material for "When Not to Kill Your Host in Competitive Bust-Boom Environments"

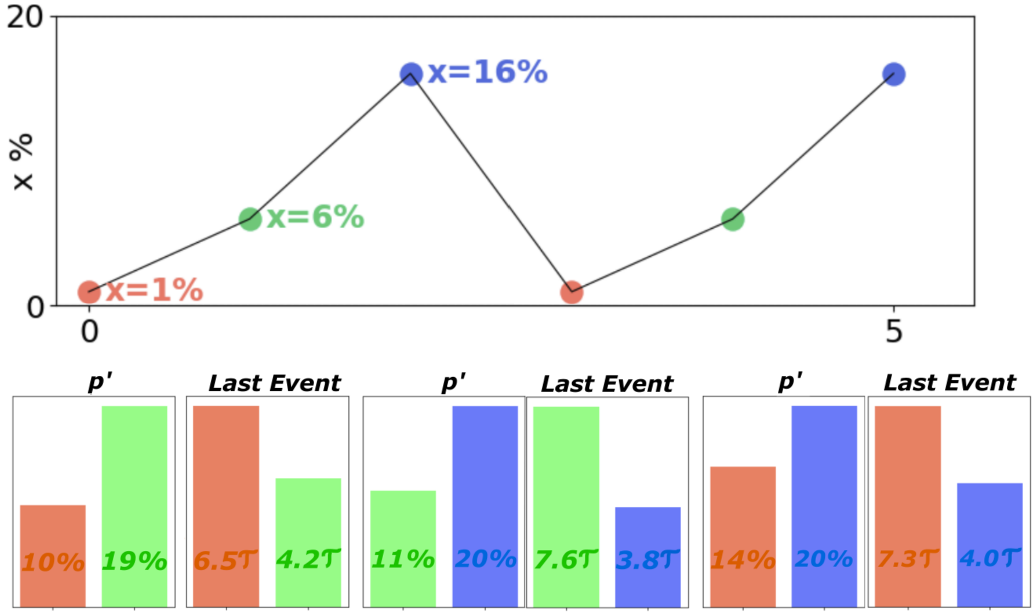


**FIG. 1.** Cyclic replacement of phages in subsequent pairwise competitions between phages without cross-immunity, with $p = 0.1, \Omega = 50$. Each dot marks the optimal strategy when competing against phage with $x$ from the previous dot, with common optimal $\sigma = 1/\tau$. The red dot marks a phage with $x = 1\%$. This is excluded by competing with a phage with $x = 6\%$ in 99.5% of time sequences where only one phages die. On these exclusions, the last 40 events contain many (∼20%) bad events, while the 0.5% of sequences favoring $x = 1\%$ the last bad event is unusually long, $t_{bad} = 6.5\tau$. Subsequently, the 6% phage is excluded by a $x = 16\%$ in 88% of sequences again characterized by many bad events (∼ 20%). Finally, the phage with $x = 16\%$ is excluded by an $x = 1\%$ phage in the 56% of sequences, typically having a last event with exceptionally long duration $t_{bad} = 7.3\tau$.

Supplementary Tables $\delta = 0$ is fixed for all simulations in main text, hence the first tabular with $\delta = 0$ is identical to presented in main text. Other tabulars increasing $\delta > 0$ corresponds to letting a fraction of free phages die before meeting a cell. Recall that the probability for a phage type, $f_x$ to meet a host is calculated by $1 - e^{-f_x/(\rho+\delta)}$. Having $\delta = 10, \Omega = 50$ corresponds to optimal $x, \sigma$ almost identical to $\delta = 0, \Omega = 5$, as first set of tables indicate. The arrows indicate cyclic replacement of optimal strategies (First cycle is explained in Fig S1).

Introducing immunity, $\sigma = 0$ is a winning strategy, since with the prophage being infinitely stable and immune, that prophage will never get excluded, therefore we copy the $\sigma_{Kelly}$ and optimize $x$. After that, we optimize $(x_A, x_{AA}, \sigma_{Kelly})$ corresponding to immune MOI game, against $(x_{optimal}, \sigma_{Kelly})$ found before. Against $(x_{optimal}, \sigma_{Kelly})$ any $(x_a < x_{optimal},\ x_{AA} > x_{optimal},\ \sigma_{Kelly})$ beats $(x_{optimal}, \sigma_{Kelly})$ 95% of the times. In summurary, in immune cases, the dose-dependant phage, is significantly stronger, when single infection lysogeny frequency is low, and when multiple infection lysogeny frequency is high.

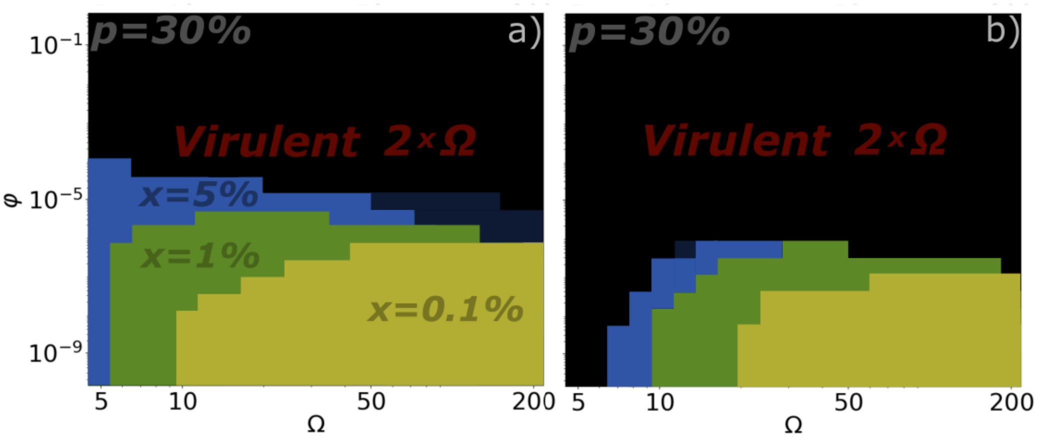


**FIG. 2.** a) shows figure from main text, with temperate phages competing against virulent phage with double $\Omega$. Only difference between a) and b), is that in a), both lysogens and phages are transfered between environments through $\varphi$. In b) only phages are transfered around. The plot indicates that in this case, also transfering lysogens, provides the temperate phage with additional bet-hedging resources.

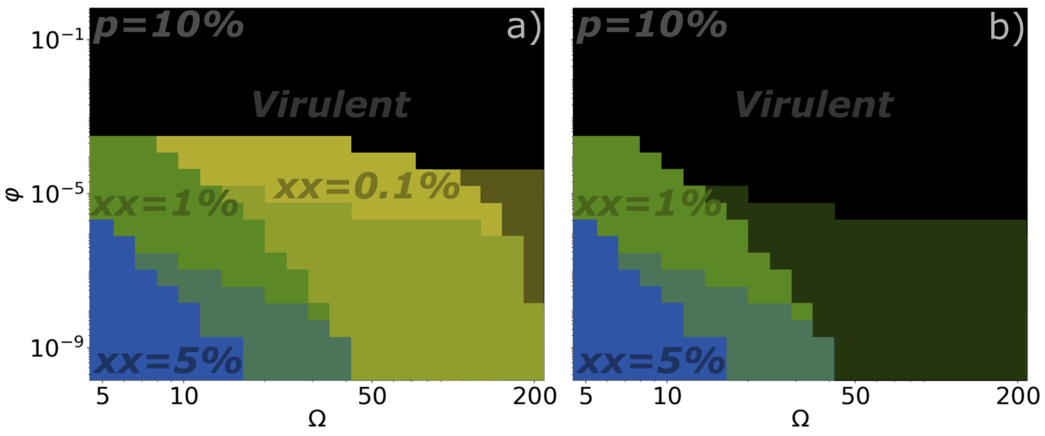


**FIG. 3.** Here dose-sensitive temperate phages are tested against virulent phage. Both the virulent and temperate phage has same burst size. For single infections, $x_A = 10^{-3}$ are fixed for all strategies. For multiple infections the lysogeny frequency is varied, either to be 1% or 5%. In panel a) $x_{AA} = x_A = 10^{-3}$ is shown, corresponding to $x = 10^{-3}$ strategy for dose-incencitive phage. In panel b), $x_A = x_{AA} = 10^{-3}$ is not shown. Key difference when $x_A = 10^{-3}$ as oppose to $x_A = 0$ is that shaded areas come back, indicating more cases where both the virulent- and the temperate phage co-exists for $10^{10}$ simulation steps.

#### I. SUPPLEMENTARY ALGORITHMS

##### A. Dose-sensitive phages

In the paper we typically considered two phages that compete for a limited pool of bacteria, meaning that growth in good periods is limited by competition for lysogens. The main model is described in the main text of the paper. Here, we consider a case where the infecting phage can change lysogeny frequency, $x$ upon multiple infections.
For separating probability for lysogenization with single $x_A$ and with multiple infection $x_{AA}$ we calculate the probability that a given bacteria is infected by at least one $A$ phage,

**TABLE I.** No cross immunity, hence with cross infections

| $p$ | $\Omega$ | $\delta$ | $(x\%, \sigma\cdot\tau)$ | $x_A\%$ | $x_{AA}\%$ |
|---|---|---|---|---|---|
| 0.1 | 5 | 0 | 2,0.8 | 1 | 2.5 |
| 0.2 | 5 | 0 | (4.5,0.3) | 4.5 | 5.5 |
| 0.3 | 5 | 0 | (7,0.1) | 6.3 | 7.5 |
| 0.1 | 50 | 0 | 1,1.0$\rightarrow$ 6,1.0 $\rightarrow$ 16,1.0 | 30 | 5 |
| 0.2 | 50 | 0 | 4,0.6 $\rightarrow$ 14,0.6 $\rightarrow$ 24,0.6 | 35 | 10 |
| 0.3 | 50 | 0 | (30 $\pm$ 20, 0.3) | 36 | 25 |
| 0.1 | 5 | 1 | 1.5,0.3$\rightarrow$1.5,0.4$\rightarrow$2,0.4 | 1.2 | 2.5 |
| 0.2 | 5 | 1 | (4,0.07) | 3.5 | 6.5 |
| 0.3 | 5 | 1 | $(10 \pm 9, 0.07 \pm 0.06)$ | 8 | 20 |
| 0.1 | 50 | 1 | 0.3,1.5$\rightarrow$2,1.2$\rightarrow$4,0.7 | 7.8 | 0.2 |
| 0.2 | 50 | 1 | $(40 \pm 38, 0.4 \pm 0.3)$ | 20 | 5 |
| 0.3 | 50 | 1 | $(30 \pm 25, 0.2 \pm 1)$ | 32 | 2 |
| 0.1 | 50 | 10 | 1.5,0.7 | 0.8 | 2 |
| 0.2 | 50 | 10 | 4.5,0.3 | 4.5 | 5 |
| 0.3 | 50 | 10 | 6,0.07 | 6 | 6.5 |

**TABLE II.** With cross immunity, hence no cross infections

| $p$ | $\Omega$ | $\delta$ | $\sigma\cdot\tau$ | $x\%$ | $x_A\%$ | $x_{AA}\%$ |
|---|---|---|---|---|---|---|
| 0.1 | 5 | 0 | 0.38 | 7.5 | [0,6] | [9,100] |
| 0.2 | 5 | 0 | 0.16 | 10.5 | [0,9.5] | [12,100] |
| 0.3 | 5 | 0 | 0.04 | 11 | [0,10] | [13,100] |
| 0.1 | 50 | 0 | 0.48 | 15 | [0,14] | [16,100] |
| 0.2 | 50 | 0 | 0.38 | 23 | [0,22] | [24,100] |
| 0.3 | 50 | 0 | 0.20 | 25.5 | [0,24] | [24,100] |
| 0.1 | 5 | 1 | 0.38 | 6 | [0, 5] | [7, 100] |
| 0.2 | 5 | 1 | 0.16 | 7 | [0,6] | [8,100] |
| 0.3 | 5 | 1 | 0.04 | 8.5 | [2,7] | [10,100] |
| 0.1 | 50 | 1 | 0.48 | 12 | [0,10] | [14,100] |
| 0.2 | 50 | 1 | 0.38 | 19 | [0,18] | [20,100] |
| 0.3 | 50 | 1 | 0.20 | 21.5 | [0,21] | [22,100] |
| 0.1 | 50 | 10 | 0.38 | 6.7 | [0,6] | [0.07,1] |
| 0.2 | 50 | 10 | 0.16 | 9.8 | [0,9] | [0.1,1] |
| 0.3 | 50 | 10 | 0.04 | 10.3 | [0,0.1] | [0.11,1] |
| 0.1 | 50 | 10 | 0.48 | 6.9 | [0,6] | [0.07,1] |
| 0.2 | 50 | 10 | 0.38 | 10.4 | [0,9] | [0.1,1] |
| 0.3 | 50 | 10 | 0.20 | 11.5 | [0,0.11] | [0.12,1] |

infected by exactly one $A$ phage, infected by more than one $A$ phage, repsictively:

$$
\begin{aligned}
F_A &= 1 - e^{-f_A/(\rho+\delta)}, \\
F_A^s &= F_A e^{-f_A/(\rho+\delta)} \\
F_A^m &= 1 - 2e^{-f_A/(\rho+\delta)} + e^{-2f_A/(\rho+\delta)}
\end{aligned}
$$

for phage-type $A$ and similarly for phage-type $B$. The replace the update during good times within eq. 5 is then replaced with:

$$\begin{aligned}
\rho_A^* &= \rho_A + x_A \cdot \rho_0 \cdot F_A^s \cdot (1 - F_B) - \rho_A \cdot F_B \\
&\quad + x_{AA} \cdot \rho_0 \cdot F_A^m \cdot (1 - F_B) \\
\rho_B^* &= \rho_B + x_B \cdot \rho_0 \cdot F_B^s \cdot (1 - F_A) - \rho_B \cdot F_A \\
&\quad + x_{BB} \cdot \rho_0 \cdot F_B^m \cdot (1 - F_A) \\
\rho_{AB}^* &= \rho_{AB} + x_A \cdot \rho_B \cdot F_A^s + x_{AA} \cdot \rho_B \cdot F_A^m \\
&\quad + x_B \cdot \rho_A \cdot F_B^s + x_{BB} \cdot \rho_A \cdot F_B^m \\
&\quad + F_A^s \cdot F_B^s \cdot \rho_0 \cdot x_A \cdot x_B \\
&\quad + F_A^m \cdot F_B^s \cdot \rho_0 \cdot x_{AA} \cdot x_B \\
&\quad + F_A^s \cdot F_B^m \cdot \rho_0 \cdot x_A \cdot x_{BB} \\
&\quad + F_A^m \cdot F_B^m \cdot \rho_0 \cdot x_{AA} \cdot x_{BB} \\
\rho_0^* &= \rho_0 \cdot (1 - F_A - F_B + F_A \cdot F_B)
\end{aligned} \tag{1}$$

$$\begin{aligned}
f_A^* &= (1 - x_A) \cdot \rho_0 \cdot F_A^s \cdot \left(1 - F_B \cdot \frac{f_B \cdot P_{lyt}(B)}{f_A + f_B}\right) \cdot \Omega \\
&\quad + (1 - x_A) \cdot \rho_B \cdot F_A^s \cdot \Omega \\
&\quad + (1 - x_{AA}) \cdot \rho_0 \cdot F_A^m \cdot \left(1 - F_B \cdot \frac{f_B \cdot P_{lyt}(B)}{f_A + f_B}\right) \cdot \Omega \\
&\quad + (1 - x_{AA}) \cdot \rho_B \cdot F_A^m \cdot \Omega \\
f_B^* &= (1 - x_B) \cdot \rho_0 \cdot F_B^s \cdot \left(1 - F_A \cdot \frac{f_A \cdot P_{lyt}(A)}{f_A + f_B}\right) \cdot \Omega \\
&\quad + (1 - x_B) \cdot \rho_A \cdot F_B^s \cdot \Omega \\
&\quad + (1 - x_{BB}) \cdot \rho_0 \cdot F_B^m \cdot \left(1 - F_A \cdot \frac{f_A \cdot P_{lyt}(A)}{f_A + f_B}\right) \cdot \Omega \\
&\quad + (1 - x_{BB}) \cdot \rho_A \cdot F_B^m \cdot \Omega \\
&\quad and\ \rho_A = \rho_A^* \ \ and\ \rho_B = \rho_B^* \ \ and\ \rho_{AB} = \rho_{AB}^* \\
&\quad and\ \rho_0 = \rho_0^* \ and\ f_A = f_A^* \ \ and\ f_B = f_B^*
\end{aligned}$$

and then continue the period with spontaneous decay from the lysogens as in the standard model. We implicitly assume that the first phage type that enters the bacterium decides the infection direction. In the above, we need probability to go lytic when infecting a bacteria, $P_{lyt}(A)$ and $P_{lyt}(B)$ that was imply $1 - x_A$ and $1 - x_B$ in eq. 5. Now it depends on single versus multiple infections:

$$\begin{aligned}
P_{lyt}(A) &= 1 - (x_A F_A^s + x_{AA} F_A^m)/F_A \\
P_{lyt}(B) &= 1 - (x_B F_A^s + x_{BB} F_B^m)/F_B
\end{aligned}$$

where we calculate the probability of lysogeny dependent on single infections and multiple infections.

If the two phages have the same immunity region, then eq. 1 is replaced by:

$$\begin{aligned}
\rho_A^* &= \rho_A + \rho_0 \cdot (x_A F_A^s + x_{AA} F_A^m) \cdot \left(1 - \frac{F_B \cdot f_B}{f_B + f_A}\right) \\
\rho_B^* &= \rho_B + \rho_0 \cdot (x_B F_B^s + x_{BB} F_B^m) \cdot \left(1 - \frac{F_A \cdot f_A}{f_B + f_A}\right) \\
\rho_0^* &= \rho_0 \cdot (1 - F_A - F_B + F_A \cdot F_B) \\
f_A^* &= ((1 - x_A) \cdot F_A^s + (1 - x_{AA}) \cdot F_A^m) \cdot \rho_0 \cdot \left(1 - \frac{f_B \cdot F_B}{f_A + f_B}\right) \cdot \Omega \\
f_B^* &= ((1 - x_B) \cdot F_B^s + (1 - x_{BB}) \cdot F_B^m) \cdot \rho_0 \cdot \left(1 - \frac{f_A \cdot F_A}{f_A + f_B}\right) \cdot \Omega \\
& \quad and\ \rho_A = \rho_A^* \ \ and\ \rho_B = \rho_B^* \ \ and\ \rho_0 = \rho_0^* \\
& \quad and\ f_A = f_A^* \ \ and\ f_B = f_B^*
\end{aligned} \tag{2}$$

which represents the generalization of eq. 5 in main to differentiate between single and double infections (remember that $F_A = F_A^s + F_A^m$).

#### B. Coupled environments

Considering coupled environments, the exchange of say virus type $f_A$ is described by the update during good periods:

$$f_A = f_A + \varphi \cdot \langle f_A \rangle - \varphi \cdot f_A \tag{3}$$

Notice $f_A = 0$ during bad periods, and the virus dies when entering a system with a bad period. Because the bad periods dominate the average $\langle f_A \rangle$ is much smaller than $f_A$:

$$\begin{aligned}
\langle f_A \rangle &= \frac{\sum_{good-events} f_A}{\sum_{good-events} 1 + \sum_{bad-events} t_{bad}} \\
&\approx \frac{1/p}{\tau + 1/p} \cdot \langle f_A(good\ times) \rangle
\end{aligned} \tag{4}$$

where the last approximation would be accurate if $f_A$ and the duration of the good period was independent. Here it is not used and only included to illustrate that $\langle f_A \rangle << \langle f_A(good\ times) \rangle$. This is because good periods are expected to be shorter than bad periods, $1/p << \tau$.

However, estimating lysogen transfering through flux, becomes more complicated, since theese are also transfered through bad periods. Following derivation, allows us to calculate average lysogeny populations, during bad periods including exchange of lysogens with other environments.

Probability for bad period $t$ to $t + dt$:

$$Bad\ period\ length:\ P(t)dt = \frac{dt}{\tau} \cdot e^{-t/\tau} \tag{5}$$

Considering flux in bad periods, giving

$$\begin{aligned}
\frac{d\rho_x}{dt} &= -(\sigma + \varphi)\rho_x + \varphi \langle \rho_x \rangle \\
\rho_x(t) &= \frac{\langle \rho_x \rangle \varphi}{\sigma + \varphi} + \left(\rho_x^0 - \frac{\langle \rho_x \rangle \varphi}{\sigma + \varphi}\right) e^{-(\sigma+\varphi)t}
\end{aligned}$$

Now call $a = \frac{\langle\rho_x\rangle\varphi}{\sigma+\varphi}$ and $b = (\rho_0 - a)$. Here, $\rho_X^0$ is a notation for lysogen density at the end of a good period. Then calculate the average $\rho_x$ given that bad period has duration $t^*$:

$$\begin{aligned}\langle\rho_x|t^*\rangle &= \frac{1}{t^*}\int_0^{t^*}\rho_x(t)dt\\ &= a + \frac{b}{t^*(\sigma+\varphi)}\left(1-e^{-(\sigma+\varphi)t^*}\right)\end{aligned}$$

Then calculate

$$\begin{aligned}\langle\rho_x\rangle &= \int_0^\infty \langle\rho_x|t^*\rangle\cdot\frac{1}{\tau}e^{-t^*/\tau}\cdot dt^*\\ &= a + \frac{b}{\tau(\sigma+\varphi)}\cdot\int_0^\infty\frac{dt^*}{t^*}e^{-t^*/\tau}\cdot\left(1-e^{-(\sigma+\varphi)t^*}\right)\\ &= a + \frac{b}{\tau(\sigma+\varphi)}\cdot\int_0^\infty dx\cdot\frac{exp(-x)-exp(-(1+\tau\sigma+\tau\varphi)x)}{x}\\ &= a + \frac{b\cdot ln(1+\tau\sigma+\tau\varphi)}{\tau(\sigma+\varphi)}\\ &= \frac{\langle\rho_x\rangle\varphi}{\sigma+\varphi} + \left(\rho_x^0 - \frac{\langle\rho_x\rangle\varphi}{\sigma+\varphi}\right)\frac{ln(1+\tau\sigma+\tau\varphi)}{\tau(\sigma+\varphi)}\end{aligned}$$

where we in large rewrite substitute $x = t^*/\tau$ and $dx = dt^*/\tau$. The last equation is a simple linear equation in one variable and can be solved for $\langle\rho_x\rangle$ given $\rho_x^0$ at the beginning of bad periods (end of good periods):

$$\langle\rho_x\rangle\left(1+\frac{\varphi}{\sigma+\varphi}\cdot(\frac{ln(1+\tau\sigma+\tau\varphi)}{\tau(\sigma+\varphi)}-1)\right) = \rho_x^0\cdot\frac{ln(1+\tau\sigma+\tau\varphi)}{\tau(\sigma+\varphi)}$$

Obtaining:

$$\langle\rho_x\rangle = \rho_x^0\cdot\frac{\ln(1+\tau\sigma+\tau\varphi)}{\left[\tau\sigma+\frac{\varphi}{\sigma+\varphi}\cdot\ln(1+\tau\sigma+\tau\varphi)\right]}.$$

which describes the mean density of lysogen type $x$ given that end of good period had density of these lysogens $\rho_x^0$. Now to use this we should estimate the average over $\rho_x^0$ over all environments. This we do by averaging over previous history:

$$\langle\langle\rho_x\rangle\rangle = \langle\rho_x^0\rangle_{previous\ 20\ good\ events}\cdot\frac{\ln(1+\tau\sigma+\tau\varphi)}{\left[\tau\sigma+\frac{\varphi}{\sigma+\varphi}\cdot\ln(1+\tau\sigma+\tau\varphi)\right]}$$

we have to start with the first good period where there no previous lysogens, and never average over more than we have until we reach 20 good periods. After this, we always average over 20 last events good period in each trajectory (also try to average over all periods in each trajectory). The update for densities after each bad period (also equal start of a good period is):

$$\rho_x = \frac{\langle\langle\rho_x\rangle\rangle\varphi}{\sigma+\varphi} + \left(\rho_x^0 - \frac{\langle\langle\rho_x\rangle\rangle\varphi}{\sigma+\varphi}\right)e^{-(\sigma+\varphi)t} \quad (6)$$

During good a period we should also allow for transfer between environments, i.e.

$$\rho_x \to \rho_x + \varphi\cdot(\langle\langle\rho_x\rangle\rangle - \rho_x) \quad (7)$$

Simulating coupled environments, by making correspondence with past events, using same update rules as usual, apart from adding average phages and lysogens and subtracting phages and lysogens. This is done by in the beginning of each good event, adding

$\varphi \cdot (\langle \rho_x \rangle + \langle f_x \rangle)$ to phage-type x, where the average is calculated over the last 500 events. 500 events is arbitrary chosen, it should be long enough to have a representable history of bad event. Simulating over anything above 250 last events had no difference in simulation outcomes. In the end of each simulation step, $\varphi \cdot f_x, \varphi \cdot \rho_x$, is subtracted from $f_x$ and $\rho_x$, respectively. Calculating averages and moving flux in and out of the system, makes the algorithm longer. Defining $\sigma_{AB} = \sigma_A + \sigma_B - \sigma_A \cdot \sigma_B$, to address decay rates for double lysogens. Initialize algorithm with $time = 0$.
The $\sum$ notations in bad events, sum over last 500 events, *time* is similar a sum over last 500 events, where the bad events accounts for most of the time. Summing over last lysogens, it is important to differentiate between good and bad events, since each time a bad event occurs, it is important to add $\tau_i$ of the average lysogen population, $\langle\langle \rho_x \rangle\rangle_i$ to averaged system. Recall that bad durations is typically $10^3$ events, while good events are 1 event, hence when summing populations, the weights are significantly different. Calculating phage average, it is not important to differentiate between event types, since adding phages in bad periods, leads to addition of zero phages. The updated algorithm, expanded with $\varphi$-correspondence with past are obtained by:

**Bad Times happening with probability p**

$$\text{Set } \tau = -\tau' \cdot \ln(\text{ran}), \quad \text{where ran} \in (0,1)$$

$$\langle \rho_A^0 \rangle_{20} = \sum_{t=-20}^{0} \rho_{A,t}^0 \cdot \frac{1}{20}, \quad \langle \rho_B^0 \rangle_{20} = \sum_{t=-20}^{0} \rho_{B,t}^0 \cdot \frac{1}{20} \quad \langle \rho_{AB}^0 \rangle_{20} = \sum_{t=-20}^{0} \rho_{AB,t}^0 \cdot \frac{1}{20}$$

$$\langle\langle \rho_A \rangle\rangle = \langle \rho_A^0 \rangle_{20} \cdot \frac{\ln(1+\tau\sigma_A+\tau\varphi)}{\left[\tau\sigma_A + \frac{\varphi}{\sigma_A+\varphi} \cdot \ln(1+\tau\sigma_A+\tau\varphi)\right]}$$

$$\langle\langle \rho_B \rangle\rangle = \langle \rho_B^0 \rangle_{20} \cdot \frac{\ln(1+\tau\sigma_B+\tau\varphi)}{\left[\tau\sigma_B + \frac{\varphi}{\sigma_B+\varphi} \cdot \ln(1+\tau\sigma_B+\tau\varphi)\right]}$$

$$\langle\langle \rho_{AB} \rangle\rangle = \langle \rho_{AB}^0 \rangle_{20} \cdot \frac{\ln(1+\tau\sigma_{AB}+\tau\varphi)}{\left[\tau\sigma_{AB} + \frac{\varphi}{\sigma_{AB}+\varphi} \cdot \ln(1+\tau\sigma_{AB}+\tau\varphi)\right]}$$

$$\rho_A^* = \frac{\langle\langle \rho_A \rangle\rangle \cdot \varphi}{\sigma_A+\varphi} + (\rho_{A_0}^0 - \frac{\langle\langle \rho_A \rangle\rangle \cdot \varphi}{\sigma_A+\varphi}) \cdot e^{-(\sigma_A+\varphi)\cdot\tau}$$

$$\rho_B^* = \frac{\langle\langle \rho_B \rangle\rangle \cdot \varphi}{\sigma_B+\varphi} + (\rho_{B_0}^0 - \frac{\langle\langle \rho_B \rangle\rangle \cdot \varphi}{\sigma_B+\varphi}) \cdot e^{-(\sigma_B+\varphi)\cdot\tau}$$

$$\rho_{AB}^* = \frac{\langle\langle \rho_{AB} \rangle\rangle \cdot \varphi}{\sigma_{AB}+\varphi} + (\rho_{(AB)_0}^0 - \frac{\langle\langle \rho_{AB} \rangle\rangle \cdot \varphi}{\sigma_{AB}+\varphi}) \cdot e^{-(\sigma_{AB}+\varphi)\cdot\tau}$$

$$\text{Set } \rho_A = \rho_A^*, \quad \rho_B = \rho_B^*, \quad \rho_{AB} = \rho_{AB}^*, \quad f_A = 0, \quad f_B = 0, \quad time \to time + \tau$$

**Good Times Chosen if Bad Not Selected:**

$$\langle \rho_A \rangle = \left( \sum_{t \in Good_{Events}} \rho_{A,t} + \sum_{t \in Bad_{Events}} \langle \langle \rho_A \rangle \rangle_t \cdot \tau_t \right) \cdot \frac{1}{time}$$

$$\langle \rho_B \rangle = \left( \sum_{t \in Good_{Events}} \rho_{B,t} + \sum_{t \in Bad_{Events}} \langle \langle \rho_B \rangle \rangle_t \cdot \tau_t \right) \cdot \frac{1}{time}$$

$$\langle \rho_{AB} \rangle = \left( \sum_{t \in Good_{Events}} \rho_{AB,t} + \sum_{t \in Bad_{Events}} \langle \langle \rho_{AB} \rangle \rangle_t \cdot \tau_t \right) \cdot \frac{1}{time}$$

$$\langle f_A \rangle = \sum_{t \in Events} f_{A,t} \cdot \frac{1}{time}$$

$$\langle f_B \rangle = \sum_{t \in Events} f_{B,t} \cdot \frac{1}{time}$$

$$f_A \to f_A + \varphi \cdot \langle f_A \rangle$$

$$f_B \to f_B + \varphi \cdot \langle f_B \rangle$$

$$\rho_A \to \rho_A + \varphi \cdot \langle \rho_A \rangle$$

$$\rho_B \to \rho_B + \varphi \cdot \langle \rho_B \rangle$$

$$\rho_{AB} \to \rho_{AB} + \varphi \cdot \langle \rho_{AB} \rangle$$

$$\rho_A^* = \rho_A \cdot (1 - \sigma_A)$$

$$\rho_B^* = \rho_B \cdot (1 - \sigma_B)$$

$$\rho_{AB}^* = (1 - \sigma_A - \sigma_B + \sigma_A \sigma_B) \cdot \rho_{AB}$$

$$f_A^* = f_A + \left( \sigma_A \rho_A + \sigma_A \cdot \left( 1 - \frac{\sigma_B^2}{\sigma_A + \sigma_B} \right) \rho_{AB} \right) \Omega$$

$$f_B^* = f_B + \left( \sigma_B \rho_B + \sigma_B \cdot \left( 1 - \frac{\sigma_A^2}{\sigma_A + \sigma_B} \right) \rho_{AB} \right) \Omega$$

$$\text{Set} \quad \rho_A = \rho_A^*, \quad \rho_B = \rho_B^*, \quad \rho_{AB} = \rho_{AB}^*, \quad f_A = f_A^*, \quad f_B = f_B^*$$

$$
\begin{aligned}
\text{Calculate} \quad & \rho, \quad F_A = 1 - e^{-f_A/(\rho+\delta)}, \quad F_B = 1 - e^{-f_B/(\rho+\delta)} \\
\rho_A^* &= \rho_A + x_A \cdot \rho_0 \cdot F_A \cdot (1 - F_B) - \rho_A \cdot F_B \\
\rho_B^* &= \rho_B + x_B \cdot \rho_0 \cdot F_B \cdot (1 - F_A) - \rho_B \cdot F_A \\
\rho_{AB}^* &= \rho_{AB} + x_A \cdot \rho_B \cdot F_A + x_B \cdot \rho_A \cdot F_B + \rho_0 \cdot F_A \cdot F_B \cdot x_A \cdot x_B \\
f_A^* &= (1 - x_A) \cdot \rho_0 \cdot F_A \cdot \left(1 - F_B \cdot \frac{(1 - x_B) f_B}{f_A + f_B}\right) \cdot \Omega \\
&\quad + (1 - x_A) \cdot \rho_B \cdot F_A \cdot \Omega \\
f_B^* &= (1 - x_B) \cdot \rho_0 \cdot F_B \cdot \left(1 - F_A \cdot \frac{(1 - x_A) f_A}{f_A + f_B}\right) \cdot \Omega \\
&\quad + (1 - x_B) \cdot \rho_A \cdot F_B \cdot \Omega \\
\rho_A^* &\to \rho_A^* \cdot (1 - \varphi) \\
\rho_B^* &\to \rho_B^* \cdot (1 - \varphi) \\
\rho_{AB}^* &\to \rho_{AB}^* \cdot (1 - \varphi) \\
f_A^* &\to f_A^* \cdot (1 - \varphi) \\
f_B^* &\to f_B^* \cdot (1 - \varphi) \\
\text{Set} \quad & \rho_A = \rho_A^*, \quad \rho_B = \rho_B^*, \quad \rho_{AB} = \rho_{AB}^*, \quad f_A = f_A^*, \quad f_B = f_B^* \\
\text{New Hosts} \quad & \rho_0 = 1 - \rho_A - \rho_B - \rho_{AB}, \quad \text{time} \to \text{time} + 1.
\end{aligned}
$$

This algorithm is tested for a temperate phage against a lytic phage, which simplifies the equations significantly. Say A is the lytic phage, then there is no $\rho_A, \rho_{AB}$. Applying same averaging method, and then using the algorithm [1], considering dose-dependent temperate phages.
